# Immunity vs Sociality: Adaptive evolution tests suggest social lifestyle exerts greater selection pressures than host-pathogen coevolution in the bees

**DOI:** 10.1101/2021.09.07.459261

**Authors:** Lauren Mee, Seth M Barribeau

## Abstract

Hosts and their parasites and pathogens are locked in antagonistic co-evolution. The genetic consequence of this can be seen in the rates of adaptive evolution in immunologically important loci in many taxa. As the risk of disease transmission increases we might also expect to see greater rates of adaptive evolution on genes of immune function. The evolution of sociality and its elaborations in insects represent enormous shift in disease transmission risk. Here, we examine whether sociality in the bees corresponds to changes in the rate of adaptive evolution in both classical canonical immune genes, and genes with putative immune functions identified from meta-analyses of honey-bee transcriptomic responses to infection. We find that measures of gene-wide adaptive evolution do not differ among canonical immune, non-canonical candidate immune, and background gene sets, but that branch-site adaptive evolution does increase with sociality regardless of gene category. Solitary species have greater rates of adaptive evolution in canonical immune genes than background genes, supporting the suggestion that social immune mechanisms may instead be the site of host-pathogen co-evolution in social species. We identify three genes with putative roles in immunity that warrant further attention (Vitellogenin *Vg*, *disks large 1 tumour suppressor*, and the uncharacterised protein *LOC100577972*). There are more gene family changes after the origin of sociality across all gene classes, with contractions occurring after the elaboration of sociality to complex eusociality. There are few genes or functions under adaptive selection that appear to be shared outside of specific lineages, suggesting that evolution of the immune system may be specific to individual species and their pathogen interactions.

**Significance:** Infectious disease drives rapid evolution of immune genes, but infection risk should be much higher in social species. To examine whether greater sociality drives faster immune system evolution we compared the rate of immune gene evolution in solitary, social, and highly eusocial bees. To account for possible novel immune genes in bees, we analysed classical immune genes alongside candidate immune genes inferred from other studies. Surprisingly, we find that solitary bees have the highest rate of immune gene evolution relative to background genes but that sociality is associated with rapid evolution across the whole genome. These findings suggest that 1) accelerated immune gene evolution is not universal, 2) immune gene evolution is moderated by sociality in that solitary species invest more into immune gene change, and 3) that social genomes are highly dynamic, which may obscure evolution at immunological loci. The types of immune genes and functions appear mostly lineage-specific, regardless of sociality, suggesting individual evolutionary his-tories exert more selection pressure than general patterns of greater pathogen exposure introduced by social living.

## 1 Introduction

Bees (Anthophila) vary widely in their social structure, from wholly solitary to advanced eusociality with distinct castes (*i.e.* honeybees and stingless bees). The advantages conferred by sociality have contributed to ants, bees, wasps and termites becoming ecologically dominant in their respective habitats (Brady et al., 2006; Cardinal and Danforth, 2011), but also carries greater risk of infectious disease. Insect societies can contain between tens to millions of genetically similar individuals packed into dense, interconnected communities which is ideal for pathogen transmission.

This increased risk of infection was presumed to exert strong selection for the maintenance and potential expansion of immune genes in highly social insects. However, the whole genome sequencing of honeybees found a surprising reduction in the number of immune genes relative to solitary model insects (Evans et al., 2006). One explanation for the depauperate individual immunity initially described by genome sequencing efforts is that social insects benefit from a suite of behavioural defenses that constitute “social immunity”, adaptations such as exclusion of the sick, allogrooming and hygienic and guarding behaviours which help prevent the spread of disease within a social group (Cremer et al., 2007; Wilson-Rich et al., 2009; Oxley et al., 2010; Dolezal and Toth, 2014; Cremer et al., 2018). These behaviours are absent in solitary insects and may have relaxed the need for extensive investment in individual protection, resulting in evolutionary losses of immune genes. However, subsequent sequencing of other Hymenoptera, including solitary species, suggest that the diminished immune repertoire described in honeybees (Evans et al., 2006) preceded the origin of sociality, rather than being a consequence of it (Barribeau et al., 2015; Kapheim et al., 2015).

An alternative explanation for the apparent reduction in immune genes is that we might simply be looking at the wrong genes. Much of what we understand about insect immunology and the genes that produce immune protection is drawn from well characterised model species, such as *Drosophila melanogaster*. While studies in model species have been instrumental in our understanding of insect immunity, Dipterans and Hymenopterans are separated by approx. 300 million years of evolution (Hennig, 1981), and it is possible that lineage-specific genes and pathways are being overlooked (Otani et al., 2016; Sackton, 2019). These novel genes may very well play important roles in immunity. The jewel wasp (*Nasonia vitrepennis*), for instance, up-regulates more wasp-specific genes upon immune challenge than genes shared with other insects (Sackton et al., 2013). A number of recent studies have used transcriptomic or microarray experiments to describe the immune responses of honeybees after infection or immune activation (Alaux et al., 2011; Richard et al., 2012; Doublet et al., 2017), and find a great number of immune-responsive genes that are not part of the classical immune suite from model species. These non-canonical, candidate immune genes and pathways highlight potentially novel immune responses that may explain the scarcity of canonical immune genes gleaned from *D. melanogaster*.

While further experimental work is needed to functionally confirm the immunological role of these non-canonical genes, we can take advantage of a feature of immune genes to infer which of these candidate genes are most likely to protect against infection. Hosts and pathogens are locked in antagonistic coevolution as any adaptation for host resistance exerts strong selection on pathogens, and *vice versa*. As a consequence, immune genes show elevated rates of adaptive evolution relative to non-immune genes (Obbard et al., 2006; Sackton et al., 2007; Obbard et al., 2009; Sackton, 2020). How social structure, and the commensurate greater risk of infection affects immune gene evolution remains unclear. While disease transmission risk is greater in high density populations, at its extreme in colonies of eusocial insects, the opportunity for social reduction of risk is also greatest. Differing patterns of adaptive evolution has been found on conserved canonical immune genes between advanced eusocial honeybees (obligately eusocial throughout life), primitively eusocial bumblebees (that have both solitary and social phases in their life history) and the wholly solitary leaf cutting bee (Barribeau et al., 2015). Here we expand on this preliminary survey by analysing the canonical and non-canonical immune responsive genes to identify whether there are comparable patterns of evolution between the classic and novel candidate immune genes across bees, and whether the patterns differ among social and solitary clades of bees.

The bees (Anthophila) are ideal for examining genomic causes of social evolution and the consequences of sociality on other genomic features, such as genes responsible for immunity, as 1) sociality evolved multiple times - there are at least three independent origins of sociality and two independent transitions to advanced eusociality (Kapheim et al., 2015; Rehan and Toth, 2015; Rehan et al., 2016); 2) all social lifestyles are extant, sometimes within the same monophyletic group; and 3) genomic resources are rapidly expanding. Here we use the whole genome sequence of 11 species of bees (Table 1) with representatives from across Anthophila with multiple social lifestyles ((Trapp et al., 2017), Figure 1). We then ask whether the patterns of molecular evolution of canonical and non-canonical immune genes vary according do social structure (eusocial, social, solitary). We examine two transitions from social to highly eusocial (Fig 1, including the honeybees *Apis mellifera* and *A. florea*, and the stingless bee *Melipona quadrifasciata*), three origins of sociality encompassing five social bees (*Lasioglossum albipes*, *Ceratina calcarata*, two bumblebees *Bombus terrestris*, *B. impatiens*, and *Eufriesea mexicana*), and three solitary branches (*Dufourea novaeangliae*, *Habropoda labriosa*, and *Megachile rotundata*).

**Figure 1:**
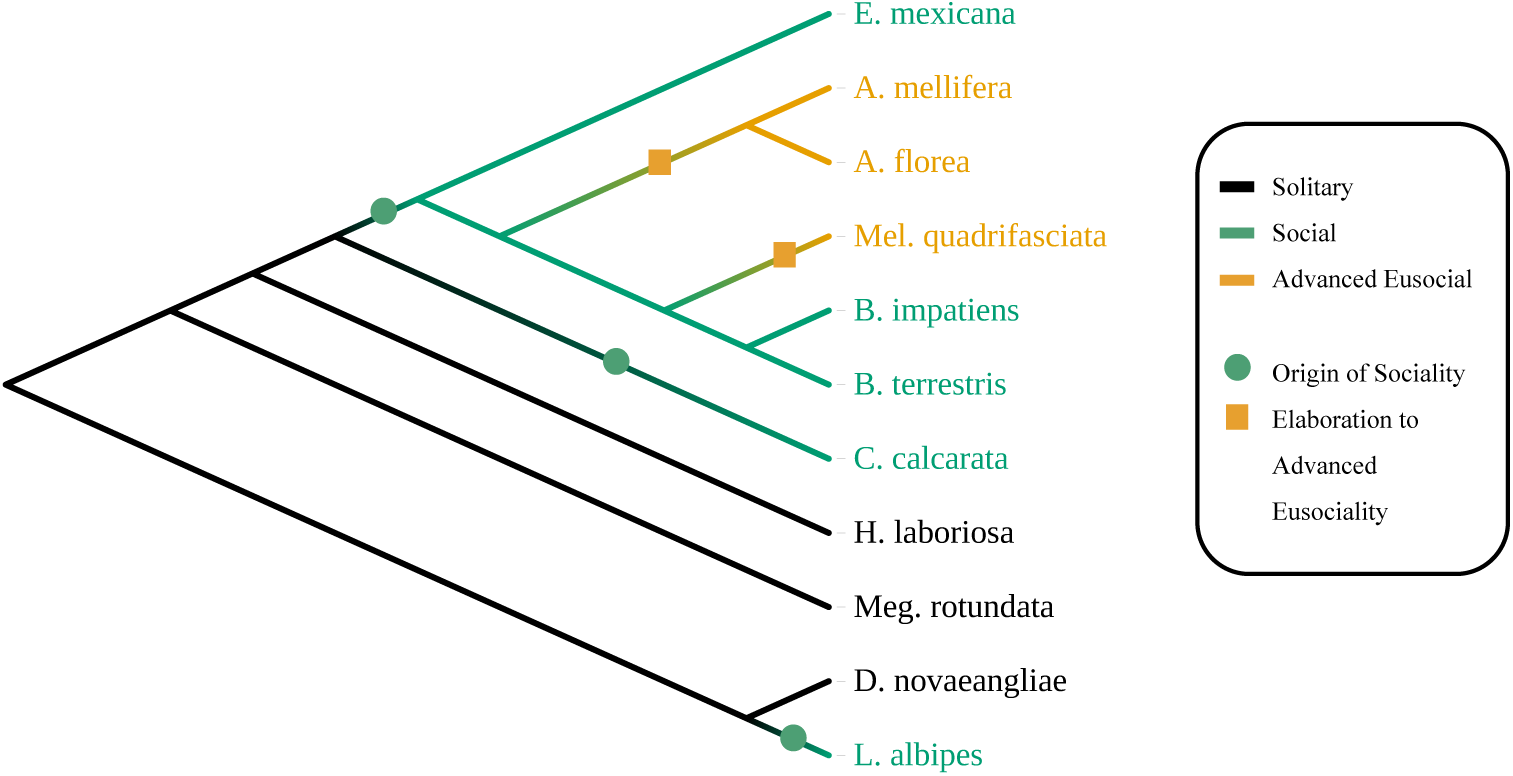
Phylogeny of the examined bee taxa. Branches are designated either solitary (black), social (green) or advanced eusocial (orange) and set as foreground in branch-specific analyses. Branch lengths are not indicative of evolutionary time.

**Table 1:** The 11 bee genome assemblies included in these analysis. Most transcriptomes and proteomes can be found on NCBI. *Melipona quadrifasciata* and *Lasioglossum albipes* are avaiable at Beebase (http://hymenopteragenome.org/beebase/)

| Family | Species | Social structure | Assembly | Reference |
| --- | --- | --- | --- | --- |
| Apidae | *Habropoda laboriosa* | Solitary | Hlab1.0 | Kapheim et al. (2015) |
|  | *Ceratina calcarata* | Social | ASM165200v1 | Rehan et al. (2016) |
|  | *Bombus terrestris* | Social | Bter1.0 | Sadd et al. (2015) |
|  | *Bombus impatiens* | Social | BIMP_2.0 | Sadd et al. (2015) |
|  | *Melipona quadrifasciata* | Complex eusocial | Mqua1.0 | Kapheim et al. (2015) |
|  | *Apis mellifera* | Complex eusocial | AmelHAv3.1 | Wallberg et al. (2019) |
|  | *Apis florea* | Complex eusocial | Aflo_1.0 | Kapheim et al. (2015) |
|  | *Eufriesea mexicana* | Social | Emex1.0 | Kapheim et al. (2015) |
| Halictidae | *Lasioglossum albipes* | Social | Lalb2.0 | Kocher et al. (2013) |
|  | *Dufourea novaeangliae* | Solitary | Dnov1.0 | Kapheim et al. (2015) |
| Megachilidae | *Megachile rotundata* | Solitary | Mrot1.0 | Kapheim et al. (2015) |

We predict that if sociality increases evolutionary pressure on immune genes then immune genes of social clades will evolve more rapidly than in solitary clades (both in adaptive sequence evolution and gene family expansion and contraction). Further, if our candidate non-canonical genes indeed serve as functional immune genes they too will be subject to the same selective pressure as canonical immune genes and demonstrate elevated rates of adaptive evolution and gene turnover relative to genes without immune function.

## 2 Materials and Methods

### 2.1 Data Resources

We used publicly available data from 11 bee species (Table 1). Whole genome sequencing derived transcriptomes and proteomes were downloaded from NCBI (NCBI Resource Coordinators, 2018) or from BeeBase (Elsik et al., 2016). We divided genes into one of three categories - canonical immune related genes, non-canonical candidate immune genes, and background genes with no known or putative immune function as a background comparison set. The canonical gene list (n = 259) was compiled from previous literature (Evans et al., 2006), the NCBI Biosystems pathway resource (Geer et al., 2009), and OrthoDB (Zdobnov et al., 2017). We collated a list of putative non-canonical immune genes (n = 1,270) from transcriptomic studies of honeybees upon immune activation through infection (Alaux et al., 2011; Richard et al., 2012; Doublet et al., 2017). Transcripts from these experimental studies were recovered from the current *A. mellifera* transcriptome build (AmelHAv3.1) and translated into their protein products. The background sample set was compiled from the rest of the *A. mellifera* genome with no known role in immunity (n = 10,594).

### 2.2 Generating Codon Alignments

We translated all genes from the *A. mellifera* genome into protein products using BLAST+ (Camacho et al., 2009). We consider these *A. mellifera* proteins as “anchor” proteins, against which other species are compared. Orthologs in the other 10 species were identified using reciprocal best-hit BLAST, keeping only those where a one-to-one ortholog was found across all species. For all genes with one-to-one orthologs across all 11 species, we produced multiple sequence protein alignments using MAFFT (Katoh et al., 2017). Each protein ortholog was reverse translated (using tBLASTn) into nucleotide sequences. These MSA/fasta pairs were then used by PAL2NAL (Suyama et al., 2006) to produce codon alignments with gaps removed for downstream PAML analyses. Alignments from non-coding transcripts were removed. By the end of this process there were 183, 872 and 4,934 alignments ready for codeml analysis in the canonical immune, non-canonical immune and background gene classes respectively.

### 2.3 Positive Selection Analyses

First, to assess the overall evolutionary rate for each gene, we used the PAML (Yang, 1997, 2007) program codeml (model=0, NSsites=0, ncatG=1) to estimate *dN/dS* ratios for each alignment. Using this metric as an indicator of rate of positive selection, we used Kruskal–Wallis one-way analyses of variance to assess differences between gene classes. To examine whether the distinct origins of sociality and social elaboration alter rate of evolution on immunologically relevant genes, we then used branch-site likelihood methods for detecting long-term shifts in positive selection in codeml (Yang and Nielsen, 2002; Zhang et al., 2005). In these models, only designated foreground branches are tested for positive selection, while all other branches in the tree are considered background. In different tests, the foreground branches were either solitary lineages, or branches following an origin or an elaboration of sociality. We consider three branches solitary, three distinct origins of sociality (one on the branch leading to the Xylocopinae, one in the corbiculate bees and one in Halictidae), and two separate elaborations to complex eusociality (one in the branch leading to the two *Apis* species and one to the stingless bees, Figure 1). We considered each lineage separately and in combination with other lineages of the same sociality (*i.e.* all social branches, all solitary branches etc.). We used the branch-site likelihood method for detecting signals of positive selection (Zhang et al., 2005). The resultant LRT statistic was compared against a *χ*^2^ distribution (*d.f* = 1), at *α* = 5%. We adjusted p-values using the Holm-Bonferonni procedure to control for multiple testing (Holm, 1979). Those with a p-value below 0.05 after correction were considered as being under positive selection.

The number of positively selected genes (PSGs) with known canonical immune and non-canonical candidate immune function were compared against the number of PSGs in the background gene class using *χ*^2^ tests. As there were two cases where PSG counts were too low to make *χ*^2^ approximation appropriate, p-values were computed by Monte Carlo simulation with 10,000 iterations using chisq.test in R (4.0.2 (R Core Team, 2020; Hope, 1968)), with Benjamini-Hochburg corrections for multiple testing (Benjamini and Hochberg, 1995). To assess trends in the proportion of genes under selection across socialities we used prop.trend.test (R 4.0.2).

### 2.4 Gene Family Analysis

Beyond adaptive evolution at coding regions, we explored whether gene families of each category grew or contracted depending on social structure. To do so, we expanded our orthology grouping to include paralogs using fastOrtho (Davis et al., 2020) on the isoform reduced proteomes used earlier. Any gene families that had less than two constituent species were dropped from the analysis. Of the resultant 8777 gene families only those that contained at least one ortholog from *A. mellifera* were analysed (n = 8283). We then used CAFE5 (Mendes et al., 2020) to statistically assess the evolutionary change in gene family number. CAFE5 uses lambda (*λ*) as a measure of the probability of both gene gain and loss (assumed equally probable) per gene per time unit across the phylogeny (De Bie et al., 2006). We calculated *λ* for each of our social structures (solitary, social, and advanced eusocial, Figure 1) on a species tree taken and modified from Rubin et al. (2019).

### 2.5 GO Analysis

We inferred gene function by assigning gene ontology terms to the isoform-reduced honeybee proteome (AmelHAv3.1) using eggNOG’s sequence mapper function set to default parameters (Huerta-Cepas et al., 2016, 2017). We then determined whether any of the gene ontology terms associated with PSGs of each gene class were significantly enriched against the complete proteome using topGO (Alexa and Rahnenfuhrer, 2016) with Fisher’s exact tests. Redundant GO terms were reduced using REVIGO (Supek et al., 2011), with similarity set to 0.5 and using the *Drosophila melanogaster* Uniprot database.

## 3 Results

### 3.1 Evolutionary Rate

In sum, we analyzed 5,989 genes for signatures of adaptive evolution. Evolutionary rate, as measured by overall gene-wide *dN/dS* ratios did not significantly differ among canonical immune genes, non-canonical immune genes, or the background set of genes (*P* = 0.1927, *df* = 2, Kruskal-Wallis one-way analysis of variance, Figure 2). One background gene alignment, *LOC724319*, gave anomalous results and was excluded from further analyses.

**Figure 2:**
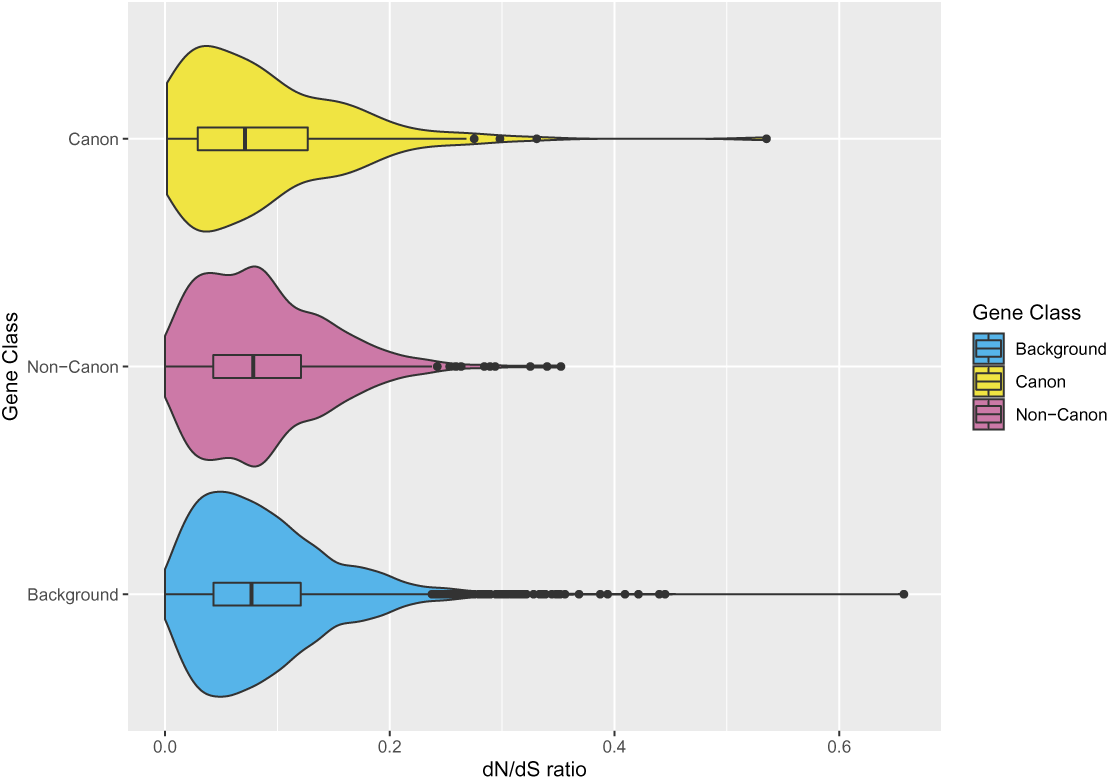
Little difference found in the evolutionary rate (dN/dS ratios) across the three gene classes: canonical immune, non-canonical candidate immune, and general background.

### 3.2 Branch-Site Models of Positive Selection

We analysed each target gene with 11 branch foregrounds (three combined sociality runs, three solitary branches, three origins of sociality, two elaborations) resulting in 65,879 tests of positive selection (see **S1-S3**) across the genome.

To assess whether the immune gene classes were under greater positive selection than the background genome, we compared the instances of canonical or non-canonical immune PSGs against those found in the background gene set (see **S4**). There were two branches where there were considerably fewer PSGs than the other lineages. These were the branches following from the origin of sociality in the corbiculates, and when all social branches were analysed as one foreground. In these tests we found no canonical immune genes under positive selection at all. There were no cases in any of the social or advanced eusocial tests of selection where the number of canonical immune PSGs were significantly greater than the background gene set. In contrast, we find more canonical immune genes under positive selection in solitary lineages (*χ*^2^ = 12.699, *P* = 0.0276). At the individual lineage level however, only the branch leading to *Habropoda laboriosa* has significantly more immune genes under positive selection (*χ*^2^ = 4.722, *P* = 0.0384), but neither of these results were significant after accounting for multiple testing. Non-canonical immune genes were not under stronger selection than the background gene set in any test, regardless of sociality (see **S4**).

We do, however, find that the overall number of PSGs increases with social complexity (Figure 3). This pattern holds regardless of the category of gene examined (canonical immune genes: *χ*^2^ = 5.1263, *df* = 1, *P* = 0.02357; non-canonical putative immune genes: *χ*^2^ = 29.625, *df* = 1, *P* =*<* 0.001; background genes: *χ*^2^ = 301.35, *df* = 2, *P* =*<* 0.001 [see **S5**]). The genes were largely sociality-specific, with the largest group of overlapping PSGs occurring between social and advanced eusocial lineages in all gene classes except canonical immune, where advanced eusocial lineages shared the same number of positively selected genes with the solitary lineages (Figure 5). Genes under selection also tend to be taxon specific with few shared genes under selection among Apidae, Halictidae, and Megachilidae (Figure 4).

**Figure 3:**
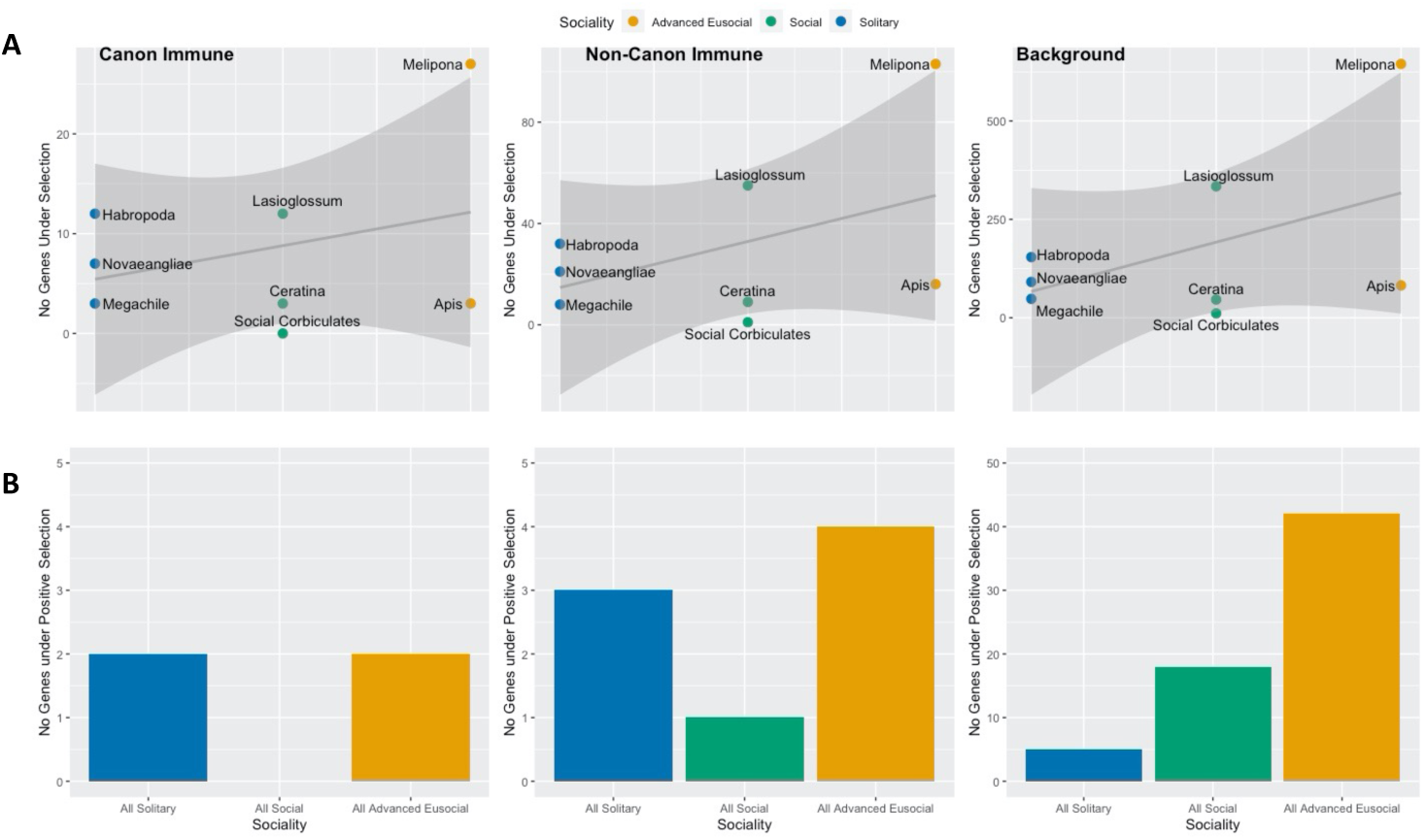
There is a linear relationship between number of genes showing signs of positive selection and how complex the social lifestyle of the bee is across all gene classes. **A**: Number of genes under positive selection plotted against sociality from solitary (blue), social (green) and advanced eusocial (orange) taxa. **B**: The number of PSGs detected per class when all branches of that degree of sociality were tested simultaneously.

**Figure 4:**
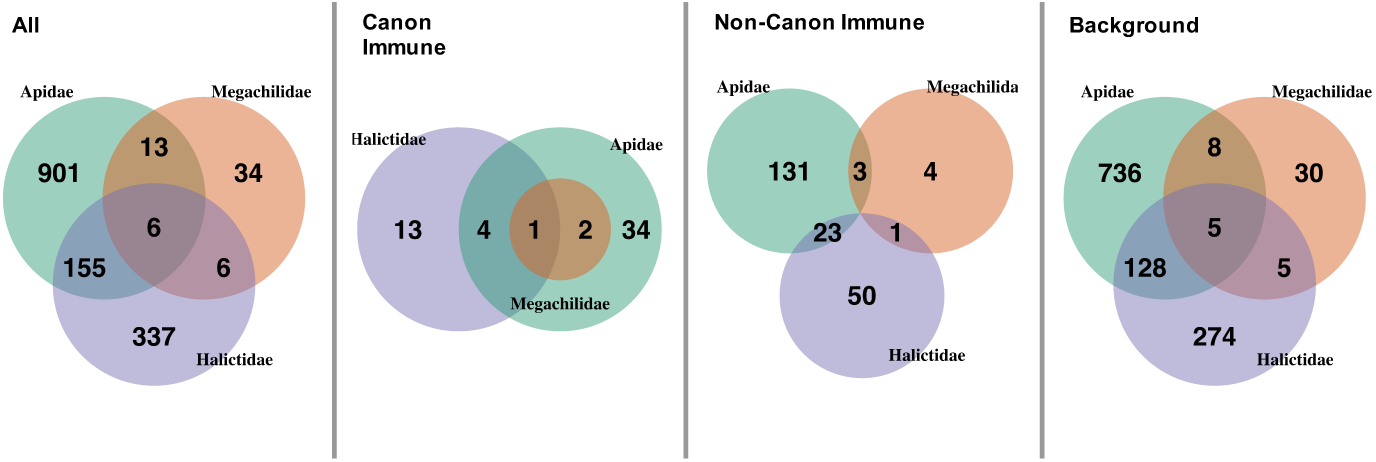
Shared genes under positive selection across the phylogeny overall, in canonical immune genes, and non-canonical candidate immune genes. There is little overlap between phylogenetic families.

**Figure 5:**
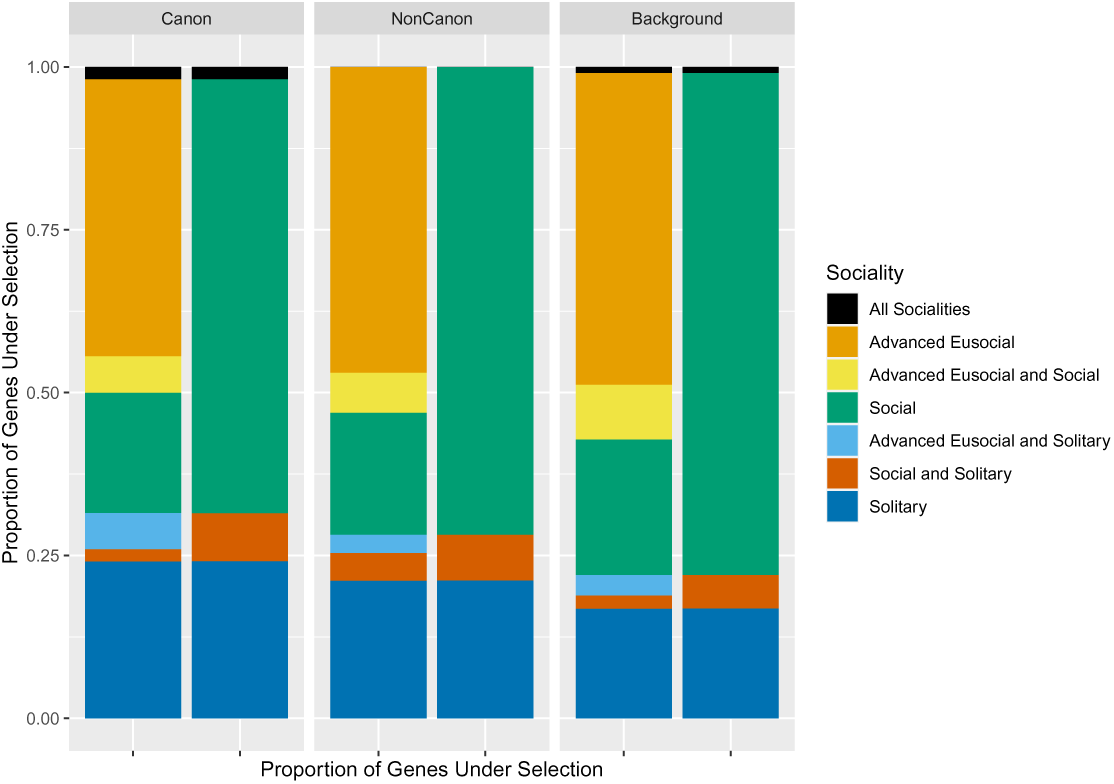
The proportion of genes under positive selection per gene class and social structure. The first bar of each graph shows all social structures and their combinations, whereas the second bar aggregates to social and solitary groups.

In total, there were 54 out of the 183 canonical immune genes that exhibit signals of positive selection (see **S1**), some of which were shared among several lineages (Table 2). *LOC726167*, a canonical immune gene, appeared the most frequently, exhibiting signs of positive selection in a social and advanced eusocial lineage (*Lasioglossum* and *Melipona*) and two solitary (*Megachile* and *Novaeangliae*) lineages. Dorsal, (*Dl*) was under selection in *Habropoda* and *Novaeangliae* and when all solitary lineages were considered simultaneously, but did not show evidence of positive selection in any social lineages.

**Table 2:** Canonical immune under positive selection (positively selected genes or PSGs) across multiple tested branches.

| Gene | Branch(es) Tested | Immune Function |
| --- | --- | --- |
| *LOC726167* | *Megachile*, *Dufourea*, *Lasioglossum*, *Melipona* | Lysosome |
| *dorsal (dl)* | All Solitary, *Habropoda*, *Dufourea* | Humoral, Toll/IMD |
| *ATG3 (LOC552315)* | *Habropoda*, All Advanced Eusocial, *Melipona* | Autophagy |
| *LOC727634* | All Solitary, *Megachile*, *Ceratina* | Autophagy |
| *dicer-1 (Dcr-1)* | *Habropoda*, *Melipona* | RNAi |
| *sialin (LOC409336)* | *Dufourea*, *Melipona* | Lysosome |
| *battenin (LOC410860)* | All Advanced, *Melipona* | Lysosome |
| *LOC411115* | *Lasioglossum*, *Apis* | Apoptosis |
| *LOC551012* | *Ceratina*, *Melipona* | Lysosome |
| *LOC551668* | *Habropoda*, *Megachile* | Autophagy |
| *LOC726947* | *Lasioglossum*, *Melipona* | Toll/IMD, Humoral |

**Table 3:** Canonical immune gene families expanded and contracted more than background and non-canonical immune genes in solitary and highly eusocial clades (Figure 6). In contrast, the background and non-canonical gene sets expanded and contracted most in the social, but not highly eusocial, clades. We denote the highest rate within social structure by italics and the highest rate of change within gene category with boldface. Overall, the highest rate of gene family expansion and contraction is in the eusocial canonical immune genes. This difference is likely driven by contractions in canonical immune gene families (Figure 6A).

| Model | $\lambda$ Solitary | $\lambda$ Social | $\lambda$ Eusocial |
| --- | --- | --- | --- |
| Background | 0.0793 | **0.1915** | 0.1645 |
| Canonical | *0.1119* | 0.1904 | ***0.2230*** |
| Non-Canonical | 0.0829 | ***0.1927*** | 0.1484 |

In the non-canonical candidate immune class there were 215 PSGs from an initial group of 872 (see **S2**). There were few non-canonical immune candidate genes under positive selection in more than 2 lineages. Vitellogenin (*Vg*) shows positive selection in the social lineages *Ceratina* and *Lasioglossum* and the solitary lineage *Novaeangliae*. An uncharacterised protein (*LOC100577972*) was a PSG in solitary *Habropoda* and the social branches leading to *Ceratina* and *Lasioglossum*. *Disks large 1 tumour suppresor protein (LOC724292)*, appears to be under positive selection in *Lasioglossum* and *Melipona* and when the social and advanced eusocial lineages are considered in combination, (see **S6**).

### 3.3 Gene Family Analysis

We find the highest probability (*λ*) of significant gene family change when considering families containing canonical immune genes in the advanced eusocial lineages (Figure 3), which this appears to be driven more by contraction in gene family number than expansion (Figure 6). Otherwise the greatest change of both non-canonical and background genes is mostly likely found in the social lineages, where there are many expansions taking place. Solitary lineages see the greatest change in the canonical immune pathway genes, but, contrary to the eusocial lineages, this appears to be due to family expansions (see **S7-9**). Of the 8283 gene families that were analysed, 224 were considered canonical immune and 1120 were classified as putatively non-canonical immune based on the *A. mellifera* orthologs present within, with all others classed as background. Of these, 32, 135 and 875 gene families (classed as canonical, non-canonical and background, respectively) were found by CAFE5 to be undergoing significant change in gene family size.

**Figure 6:**
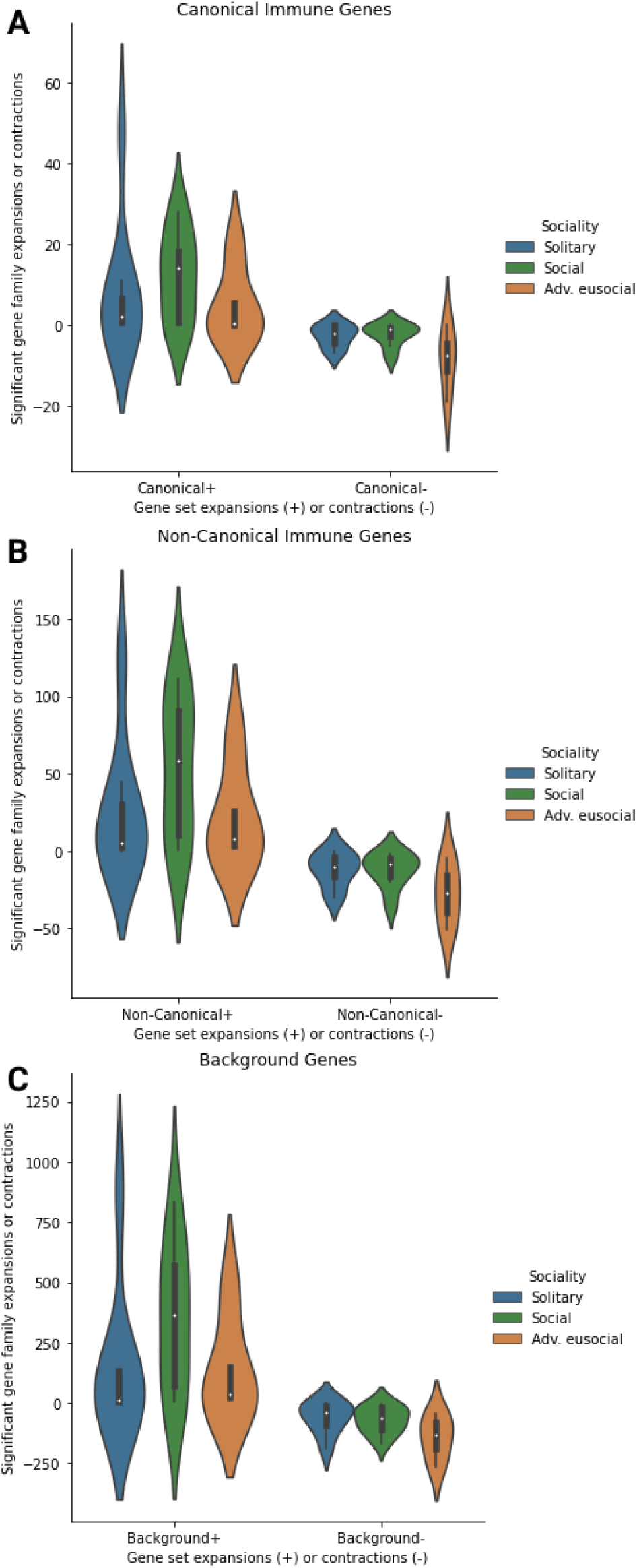
Highly eusocial clades have higher rates of signficant (A) canonical immune gene family expansions (+) and contractions (-) than less social and solitary clades, while social clades have more gene family expansions and contractions than solitary or highly eusocial clades in (B) non-canonical or (C) back-ground gene sets. Y axis denotes the number of paralogs gained or lost per family. Only significantly changing gene families are considered.

### 3.4 GO Analysis

A number of PSGs from all gene classes have enriched GO terms that are likely to be indicative of immune function. As would be expected, the class with the highest proportion of immune-associated GO terms were found in the canonical immune genes under selection regardless of sociality (Figure 7). Across the three social lifestyles, there were more immune GO terms found enriched in the non-canonical candidate PSGs than in the background, except in the social clades where the opposite was true. In the case of the advanced eusocial clades, PSGs had considerably more immune-associated GO terms in the canonical and non-canonical class of PSGs than background.

**Figure 7:**
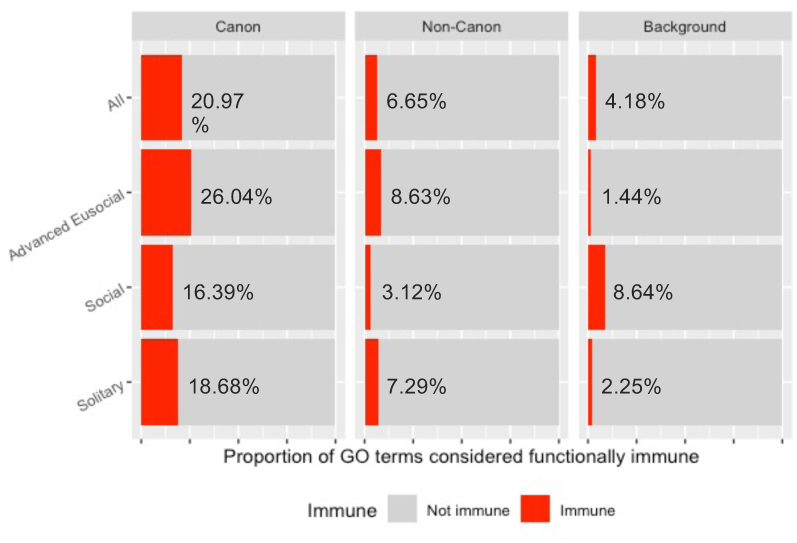
Percent of enriched immune GO terms in the PSG of each class according to sociality

When comparing across social structures, the majority of GO terms were unique to the combination of sociality and class with no enriched GO terms shared amongst all three social designations within a single class of gene (Figure 8). The largest overlap of GO terms were between social and advanced eusocial non-canonical PSGs (n = 24), but none of these GO terms have a clear association with immune function and were broadly developmental (**S10-S12**).

**Figure 8:**
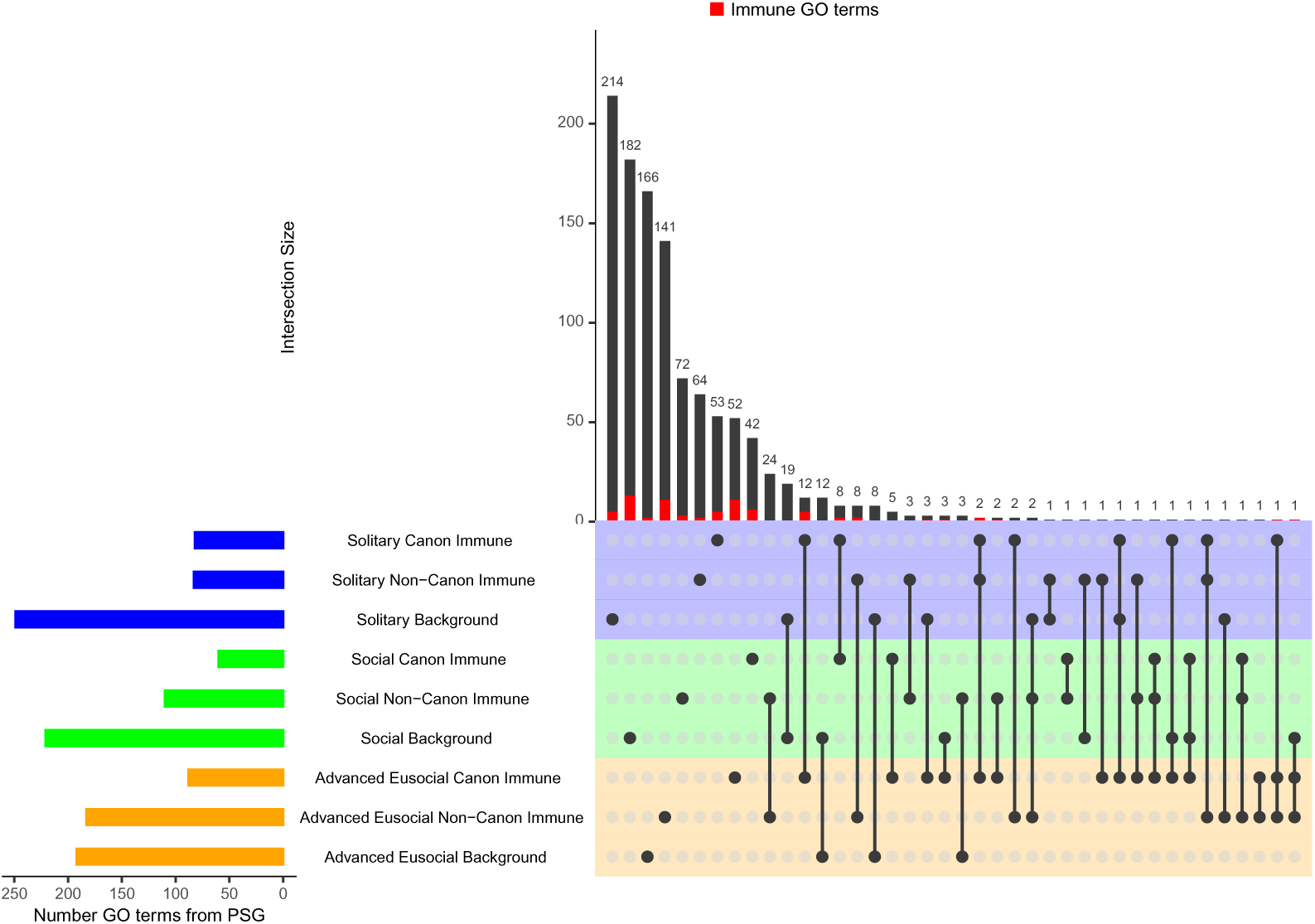
Though there are some shared enriched GO terms between the non-canonical immune PSGs of the social (highlighted in green) and advanced eusocial (highlighted in orange) lineages, there is otherwise little overlap between enriched GO terms of PSGs. There are more immune GO terms associated with the canonical and non-canonical immune PSGs than those deemed background in the solitary (highlighted in blue) and advanced eusocial lineages. GO terms that are considered immune are coloured red.

## 4 Discussion

It is near dogma that genes important for immunity evolve more rapidly than other genes. This is intuitive given the requisitely antagonistic coevolution between hosts and pathogens generally, and in genes that affect infection and immunity specifically (Sackton, 2019). However, here we find no difference between the gene-wide evolutionary rate of either known, a.k.a. canonical, immune genes, or putative non-canonical immune genes, and background genes of no suspected immune function across 11 bee species (Figure 2).

While immune genes, either established or novel, may not show faster gene-wide adaptive evolution, a more nuanced picture of the relationship between social structure and immunity begins to emerge when we examine the degree of social complexity. There appears to be more positive selection in canonical immune genes relative to the background genes of the solitary lineages, both overall, and in the branch leading to *Habropoda*. In contrast, social lineages do not show greater positive selection in canonical immune genes. This supports the suggestion that sociality affords less reliance on canonical immune genes (Evans and Armstrong, 2006). Thus, as solitary bees invest more in individual immunity, we see greater evidence of adaptive molecular evolution at known immunological loci. However, the actual proportion of canonical immune genes showing signs of selection was the same in both the all-solitary and the all-advanced branch tests (0.011, see **S4**). This reflects the much higher number of PSGs in the background genes of the social branches. It is possible that while immune genes evolve more quickly than other genes, the shift to highly social lifestyle is associated with rapid evolution of many non-immune genes (*e.g.*, genes involved in longevity, reproduction (Sadd et al., 2005; Kapheim et al., 2015; Shell et al., 2021)), thus swamping the signal of immune gene evolution that has been so well described in diverse solitary species, and which we find in our solitary bee species.

After looking at the number of PSGs across gene classes per lineage, we then collapsed those lineages into tests of the three social designations (solitary, social, highly eusocial). We detect increasing levels of selection upon the immune genes with increased social living, which may be indicative of an increase in pathogen risk leading to stronger selection (Figure 3, **S5**). As this pattern is also seen in the other gene sets, including background, it remains unclear as to whether the non-canonical immune genes are evolving due to a role in immunity or because they play a part in other processes important for social living. Changes in the immune genes of the social and advanced eusocial lineages may be harder to detect against a background genome that is transitioning between social lifestyles.

We proposed that if the putative immune genes identified from transcriptomic studies had immune functions then these genes ought to be more commonly under positive selection in social than solitary lineages. While we do see this pattern when examining all social groups together, we do not necessarily recover the same genes when examining each branch to sociality individually. For example, both the branches leading to the solitary lineages of *Habropoda* and *Dufourea* had a higher proportion of non-canonical candidate immune PSGs than the branch leading to the *Apis* species. This could still support the idea that these genes are putatively Hymenopteran-specific immune genes, regardless of sociality, or that these genes are important for other non-immune functions in solitary bees.

We find support for putative immune function of the non-canonical gene set in our GO term enrichment tests (see **S10-S12**). When tested together, the advanced eusocial branches had non-canonical PSGs enriched for functions associated with lysosomes and autophagosomes, required for phagocytyosis and autophagy (Hillyer, 2016; Kuo et al., 2018). On the branch leading to the stingless bees there were terms associated with autophagosomes, DNA repair and the recognition of peptidoglycan proteins, a component of pathogen cell walls (Wang et al., 2019). In the honeybees, there were multiple terms associated with DNA repair, which cross-talks extensively with immunity, particularly in double-stranded breaks and viral infection (Ishii et al., 2006; Xu, 2006; Brzostek-Racine et al., 2011; Z^̌^gur-Bertok, 2013). All social branches had a term involved in the regulation and signalling of the JAK/STAT pathway, a conserved component of the insect immune response (McMenamin et al., 2018; Bang, 2019), and repair of double-stranded DNA breaks enriched in the *Ceratina* and *Lasioglossum* branches. In the branch leading to *Habropoda*, we find response to dsRNA and generation of the RISC complex (including the RNA interference effector complex) were enriched, suggesting non-canonical PSGs in this lineage may play a role in responding to viral threats (Ding, 2010). Genes involved in the RNA interference (RNAi) pathway have previously been found to evolve rapidly (Obbard et al., 2006; Barribeau et al., 2015). In contrast, there were fewer immune terms in the background GO enrichment tests in solitary and advanced eusocial bees (Figure 7).

To infer more general patterns of positive selection in immune and putatively immune genes, we explored which genes were under selection in several lineages. In the canonical immune genes, *LOC726167* appeared in four lineages, two social and two solitary. *LOC726167* codes for a lipopolysaccharide-induced tumour necrosis factor-alpha factor (LITAF) homolog. In mammals, LITAF is well-characterised as a transcription factor that plays an important role in inflammation, and probably directs proteins to lysosomes for degradation (Tang et al., 2005; Zou et al., 2015). In aquatic molluscs, LITAF is involved in the immune response to bacterial challenge (Park et al., 2008; De Zoysa et al., 2010; Wang et al., 2012; Liu et al., 2020). LITAF has not been functionally tested in *D. melanogaster* or bees, but a LITAF-like protein was found to play a role in anti-parasite defense in *Anopheles gambiae* (Smith et al., 2012). The other canonical immune gene which was under selection in more than one lineage, dorsal (*dl*), has long been known to play a role in the innate immune response of insects, having been linked to anti-microbial, -fungal and -viral responses through activation of the Toll pathway (Belvin and Anderson, 1996; Silverman and Maniatis, 2001; Bangham et al., 2006; Ferreira et al., 2014; Sheehan et al., 2018). In Hymenoptera specifically, *dl* orthologs have been found to regulate expression of defensin (Loureņco et al., 2018). *Dl* also plays a role in development, and so it cannot be said with absolute certainty that the pressure behind the positive selection in the *Habropoda* and *Novaeangliae* lineages were down to adaptive immune function.

In the non-canonical candidate immune genes, vitellogenin (*Vg*) is under selection in several lineages (social: *Lasioglossum* and *Ceratina*, solitary: *Habropoda*). *Vg* is an ancient yolk precursor protein found throughout the Metazoans and functions as a large lipid transfer protein important in a number of functions, including immune responses (Hayward et al., 2010; Salmela and Sundström, 2018). *Vg* protects against oxidative stress (Seehuus et al., 2006; Havukainen et al., 2013) and is involved in trans-generational immune priming (Salmela et al., 2015; Harwood et al., 2019), where pathogen resistance is transferred from adults to offspring (Milutinovíc et al., 2016). *Vg* is also down-regulated upon *Varroa* mite or viral infection (Loureņco et al., 2009; Alaux et al., 2011; Doublet et al., 2017), perhaps as resources are reallocated from storage to immune defences. Behaviourally, down-regulation of *Vg* in tandem with up-regulation of *malvolio* results in workers switching from colony-based work to foraging, effectively removing the infection from the heart of the colony. Shifting worker tasks to minimize risk within the colony serves as a demographic and behavioural defensive adaptation which can be considered a component of social immunity (Ben-Shahar et al., 2004; Antonio et al., 2008). More directly, *Vg* can also recognise and bind pathogens (Zhang et al., 2011), suppress microbial growth, and is up-regulated in response to bacterial challenge in *Apis cerana* (Park et al., 2018). *Vg* functions in immunological and non-immune processes. Thus, while we do see adaptive evolution of *Vg* in our tests, discerning the relative role of immune or other pressures in driving these coding changes remain unclear.

*LOC100577972* is an uncharacterised protein under positive selection in the same lineages as *Vg*. There appears to be very similar homologs in all sequenced Hymenopterans, with more distant orthologs in other holometabolous species (NCBI Resource Coordinators, 2018). However, the function of these proteins has not yet been characterised and any guesses at function would be speculative. The final candidate immune gene of particular interest is *LOC724292*, or *disks large 1 tumor suppressor protein* (*dlg1*). This is the only multiple-lineage PSG found only in social branches (specifically, when all complex eusocial or social branches are considered together, and in the branches leading to stingless bees (*Melipona*) and *Lasioglossum*). *dlg1* is a membrane associated guanylate kinase (MAGUK) protein essential for neurogenesis, cell polarity and growth control in *D. melanogaster* (Woods and Bryant, 1991; Bilder et al., 2000; Mendoza et al., 2003). In *D. melanogaster dlg1* protein is important for maintaining gut epithelial integrity (Clark et al., 2015). *dlg1* is further thought to modulate innate immune signalling, as down-regulation of *dlg1* in *D. melanogaster* leads to increased expression of the antimicrobial peptide Diptericin (Xiong et al., 2016). It is similarly down-regulated in the presence of parasitoid wasps where it putatively plays a role in an anti-macroparasite response (Schlenke et al., 2007). Supporting this, *dlg1* was down-regulated in honeybees infected with *Varroa destructor* mites (Navajas et al., 2008). However, this pattern is not universal. When infected with virus, *dlg1* expression was increased in both ants and honey-bees (Galbraith et al., 2015; Manfredini et al., 2016). Whether this is due to differences in host responses to virus or viral manipulation is unclear. Across different infection studies using different pathogens and pathogen combinations, *dlg1* was neither uniformly up- or down-regulated in challenged honeybees (Doublet et al., 2017).

*dlg1* ’s putative role in immunity in insects has some support, but its role in neurogenesis may also explain positive selection in the social branches. It is often posited that sociality, and the complex interactions that sociality requires, can drive the evolution of larger or more complex brains (Dunbar, 1998). Although the complex structure of the honeybee brain relative to other insects initially supported this hypothesis in invertebrates, it rather appears that complex neurology predates sociality within Hymenoptera, as the brain structure of solitary and social Hymenopterans is comparable (Withers et al., 2008; Farris and Schulmeister, 2011; Farris, 2013). As this propensity for complexity existed in bees before sociality, selection on *dlg1* may also be due to other factors that social living introduces, such as the increased danger of pathogen transmission between nest-mates.

Beyond sequence level adaptive evolution, we further explored whether gene turnover - the expansions or contractions in the number of genes within a gene family - differ among our gene categories and among social structures. We find that social species have more gene expansions than either solitary species or advanced eusocial species (Figure 6). This is likely due to the host of new behaviours and adaptations necessary to facilitate a transition from solitary to social living. This is most pronounced in non-canonical immune genes, but we cannot say whether this is due to putative immune function or other developmental processes involved in living in insect societies.

Strikingly, advanced eusocial clades have more gene family contractions than social or solitary clades. The number of expansions within these clades is greatest in the non-canonical immune genes and background gene sets (which recapitulates the findings of Kapheim et al. (2015)). It is noteworthy that the pattern of increased gene family number in social taxa disappears in advanced eusocial clades, where instead, we see contractions in gene families, most dramatically in the canonical immune genes. This may be that as complex eusociality introduces an evolutionary “point of no return” (Wilson and Hölldobler, 2005), the continual loss of social plasticity translates to gene loss in particular families. It may well be that the presence of social immunity does indeed lead to a reduced arsenal of immune genes in the advanced eusocial insects, as other genes and pathways are utilised instead. These analyses suggests that in bees, the core canonical immune genes do not vary much in either sequence evolution or gene family membership relative to background genes, but the non-canonical genes may be less constrained.

In terms of gene function, we find that there are more immune GO terms associated with putative non-canonical immune PSGs than background in the solitary and advanced eusocial clades and when all PSGs are considered regardless of social lifestyle (Figure 7). This lends some weight to these genes being used in immune responses that are yet to be clearly elucidated. Despite this, we find relatively little overlap of function across the different social lifestyles and rather function - and PSGs - seem mostly taxon-specific (Figures 4, 8). With the use of RNAseq and immune challenge transcriptomic experiments becoming more common in the insects, a complicated picture is emerging where immune responses are challenge-specific, recruiting genes existing outside of what is considered canonical in individual host-pathogen couplings (Doublet et al., 2017; Troha et al., 2018; Sackton, 2019). This means that individual species may invest resources into very specialised pathways dependent on the prevalence of certain pathogenic species or challenges in their unique ecological niches throughout evolutionary time. Thus, it may be difficult to detect such patterns of adaptive selection at a broad phylogenetic and geographic scale.

We predicted that increased sociality increases infectious disease risk and would thus drive the rate of adaptive evolution of the genes that encode the immune system writ large; including both the classical definition of the immune response, and candidate physiological, demographic, or behavioural components. We find instead a complex pattern where the evolution of classical physiological components of immunity are accelerated in solitary bees, but that genomes in general are evolving more rapidly in the more social and advanced eusocial lineages. We interpret this as the result of solitary bees needing to invest more into individual immunity, whereas social animals have much more dynamic patterns of selection in areas other than immune as they transition into different social lifestyles. We present evidence as to some candidate genes that warrant further experimental investigation, and suggest that future work hones in on specific host-pathogen challenges in closely related populations in order to unravel the genetic underpinning of Hymenopteran immunity.

## 5 Article types

### Author Contributions

SMB conceived of the initial design. LM and SMB designed the specific tests. LM largely conducted the analyses, with some direction and occasional contributions from SMB. LM and SMB dragged one another, kicking and screaming, through various parts of this work; despite this, both largely still get along. Both LM and SMB wrote the manuscript.

### Funding

This work was supported by a NERC PhD fellowship.

## Acknowledgments

We thank Jon Bollback and the University of Liverpool Centre for Genomic Research for use of their servers for these analyses.

## Data Availability Statement

All data analyzed in this manuscript can be found publicly (Table 1). Our code, input trees and pipelines can all be found on our github repository: https://github.com/LMee17/Proj0_Analysis.

## References

C. Alaux, C. Dantec, H. Parrinello, and Y. Le Conte. Nutrigenomics in honey bees: Digital gene expression analysis of pollen’s nutritive effects on healthy and varroa-parasitized bees. BMC Genomics, 12(October), 2011. ISSN 14712164. doi: 10.1186/1471-2164-12-496.

A. Alexa and J. Rahnenfuhrer. Gene set enrichment analysis with topGO. R package version 2.30.0., 2016.

D. S. M. Antonio, K. R. Guidugli-Lazzarini, A. M. Do Nascimento, Z. L. P. Simões, and K. Hartfelder. Rnai-mediated silencing of vitellogenin gene function turns honeybee (apis mellifera) workers into extremely precocious foragers. Naturwissenschaften, 95(10):953–961, 2008.

I. S. Bang. Jak/stat signaling in insect innate immunity. Entomo- logical Research, 49(8):339–353, 2019. doi: https://doi.org/10.1111/1748-5967.12384. URL https://onlinelibrary.wiley.com/doi/abs/10.1111/1748-5967.12384.

J. Bangham, F. Jiggins, and B. Lemaitre. Insect immunity: the post-genomic era. Immunity, 25(1):1–5, 2006.

S. M. Barribeau, B. M. Sadd, L. du Plessis, M. J. Brown, S. D. Buechel, K. Cappelle, J. C. Carolan, O. Christiaens, T. J. Colgan, S. Erler, J. Evans, S. Helbing, E. Karaus, H. M. G. Lattorff, M. Marxer, I. Meeus, K. Näpflin, J. Niu, R. Schmid-Hempel, G. Smagghe, R. M. Waterhouse, N. Yu, E. M. Zdobnov, and P. Schmid-Hempel. A depauperate immune repertoire precedes evolution of sociality in bees. Genome Biology, 16 (1):83, 2015. ISSN 1465-6906. doi: 10.1186/s13059-015-0628-y. URL http://genomebiology.com/2015/16/1/83.

M. P. Belvin and K. V. Anderson. A conserved signaling pathway: the drosophila toll-dorsal pathway. Annual review of cell and developmental biology, 12(1):393–416, 1996.

Y. Ben-Shahar, N. L. Dudek, and G. E. Robinson. Phenotypic deconstruction reveals involvement of manganese transporter malvolio in honey bee division of labor. Journal of experimental biology, 207(19):3281–3288, 2004.

Y. Benjamini and Y. Hochberg. Controlling the False Discovery Rate : A Practical and Powerful Approach to Multiple Testing. Journal of the Royal Statistical Society. Series B ( Methodological ), 57(1):289–300, 1995.

D. Bilder, M. Li, and N. Perrimon. Cooperative regulation of cell polarity and growth by drosophila tumor suppressors. Science, 289(5476):113–116, 2000.

S. G. Brady, S. Sipes, A. Pearson, and B. N. Danforth. Recent and simultaneous origins of eusociality in halictid bees. Proceedings of the Royal Society B: Biological Sciences, 273(1594):1643–1649, 2006. ISSN 0962-8452. doi: 10.1098/rspb.2006.3496. URL http://rspb.royalsocietypublishing.org/cgi/doi/10.1098/rspb.2006.3496.

S. Brzostek-Racine, C. Gordon, S. Van Scoy, and N. C. Reich. The dna damage response induces ifn. Journal of immunology (Baltimore, Md. : 1950), 187(10):5336–5345, 11 2011. doi: 10.4049/jimmunol.1100040. URL https://pubmed.ncbi.nlm.nih.gov/22013119.

C. Camacho, G. Coulouris, V. Avagyan, N. Ma, J. Papadopoulos, K. Bealer, and T. L. Madden. Blast+: architecture and applications. BMC bioinformatics, 10(1):1–9, 2009.

S. Cardinal and B. N. Danforth. The antiquity and evolutionary history of social behavior in bees. PLoS ONE, 6(6), 2011. ISSN 19326203. doi: 10.1371/journal.pone.0021086.

R. I. Clark, A. Salazar, R. Yamada, S. Fitz-Gibbon, M. Morselli, J. Alcaraz, A. Rana, M. Rera, M. Pellegrini, W. J. William, et al. Distinct shifts in microbiota composition during drosophila aging impair intestinal function and drive mortality. Cell reports, 12(10):1656–1667, 2015.

S. Cremer, S. A. Armitage, and P. Schmid-Hempel. Social Immunity. Current Biology, 17(16):693–702, 2007. ISSN 09609822. doi: 10.1016/j.cub.2007.06.008.

S. Cremer, C. D. Pull, and M. A. Fürst. Social Immunity: Emergence and Evolution of Colony-Level Disease Protection. Annual Review of Entomology, 63:105–123, 2018. ISSN 0066-4170. doi: 10.1146/annurev-ento-020117-043110. URL http://www.annualreviews.org/doi/10.1146/annurev-ento-020117-043110.

J. J. Davis, A. R. Wattam, R. K. Aziz, T. Brettin, R. Butler, R. M. Butler, P. Chlenski, N. Conrad, A. Dickerman, E. M. Dietrich, J. L. Gabbard, S. Gerdes, A. Guard, R. W. Kenyon, D. Machi, C. Mao, D. Murphy-Olson, M. Nguyen, E. K. Nordberg, G. J. Olsen, R. D. Olson, J. C. Overbeek, R. Overbeek, B. Parrello, G. D. Pusch, M. Shukla, C. Thomas, M. VanOeffelen, V. Vonstein, A. S. Warren, F. Xia, D. Xie, H. Yoo, and R. Stevens. The PATRIC bioinformatics resource center: expanding data and analysis capabilities. Nucleic Acids Res., 48(D1):D606–D612, Jan. 2020.

T. De Bie, N. Cristianini, J. P. Demuth, and M. W. Hahn. Cafe: a computational tool for the study of gene family evolution. Bioinformatics, 22(10): 1269–1271, 2006.

M. De Zoysa, C. Nikapitiya, C. Oh, I. Whang, J.-S. Lee, S.-J. Jung, C. Y. Choi, and J. Lee. Molecular evidence for the existence of lipopolysaccharide- induced tnf-*α* factor (litaf) and rel/nf-kb pathways in disk abalone (haliotis discus discus). Fish & Shellfish Immunology, 28(5):754–763, 2010. ISSN 1050-4648. doi: https://doi.org/10.1016/j.fsi.2010.01.024. URL https://www.sciencedirect.com/science/article/pii/S1050464810000379.

S.-W. Ding. Rna-based antiviral immunity. Nature Reviews Immunology, 10(9):632–644, 2010. doi: 10.1038/nri2824. URL https://doi.org/10.1038/nri2824.

A. G. Dolezal and A. L. Toth. Honey bee sociogenomics: A genome-scale perspective on bee social behavior and health. Apidologie, 45(3):375–395, 2014. ISSN 12979678. doi: 10.1007/s13592-013-0251-4.

V. Doublet, Y. Poeschl, A. Gogol-Döring, C. Alaux, D. Annoscia, C. Aurori, S. M. Barribeau, O. C. Bedoya-Reina, M. J. F. Brown, J. C. Bull, M. L. Flenniken, D. A. Galbraith, E. Genersch, S. Gisder, I. Grosse, H. L. Holt, D. Hultmark, H. M. G. Lattorff, Y. Le Conte, F. Manfredini, D. P. McMahon, R. F. A. Moritz, F. Nazzi, E. L. Ninõ, K. Nowick, R. P. van Rij, R. J. Paxton, and C. M. Grozinger. Unity in de- fence: honeybee workers exhibit conserved molecular responses to diverse pathogens. BMC Genomics, 18:207, 2017. ISSN 1471-2164. doi: 10.1186/s12864-017-3624-7. URL http://bmcgenomics.biomedcentral.com/articles/10.1186/s12864-017-3624-7.

R. I. Dunbar. The social brain hypothesis. Evolutionary Anthropology: Issues, News, and Reviews: Issues, News, and Reviews, 6(5):178–190, 1998.

C. G. Elsik, A. Tayal, C. M. Diesh, D. R. Unni, M. L. Emery, H. N. Nguyen, and D. E. Hagen. Hymenoptera Genome Database: Integrating genome annotations in HymenopteraMine. Nucleic Acids Research, 44(D1):D793– D800, 2016. ISSN 13624962. doi: 10.1093/nar/gkv1208.

J. D. Evans and T.-N. Armstrong. Antagonistic interactions between honey bee bacterial symbionts and implications for disease. BMC ecology, 6:4, 2006. ISSN 1472-6785. doi: 10.1186/1472-6785-6-4. URL http://www.ncbi.nlm.nih.gov/pubmed/16551367.

J. D. Evans, K. Aronstein, Y. P. Chen, C. Hetru, J.-L. L. Imler, H. Jiang, M. Kanost, G. J. Thompson, Z. Zou, and D. Hult- mark. Immune pathways and defence mechanisms in honey bees Apis mellifera. Insect Mol Biol, 15(5):645–656, 2006. ISSN 0962-1075. doi: 10.1111/j.1365-2583.2006.00682.x. URL http://www.pubmedcentral.nih.gov/articlerender.fcgi?artid=1847501{&}tool=pmcentrez{&}rendertype=abstract{%}5Cnhttp://www.ncbi.nlm.nih.gov/entrez/query.fcgi?cmd=Retrieve{&}db=PubMed{&}dopt=Citation{&}list{_}uids=17069638.

S. M. Farris. Evolution of complex higher brain centers and behaviors: behavioral correlates of mushroom body elaboration in insects. Brain, behavior and evolution, 82(1):9–18, 2013.

S. M. Farris and S. Schulmeister. Parasitoidism, not sociality, is associated with the evolution of elaborate mushroom bodies in the brains of hymenopteran insects. Proceedings of the Royal Society B: Biological Sciences, 278(1707): 940–951, 2011.

Á. G. Ferreira, H. Naylor, S. S. Esteves, I. S. Pais, N. E. Martins, and L. Teixeira. The toll-dorsal pathway is required for resistance to viral oral infection in drosophila. PLoS pathogens, 10(12):e1004507, 2014.

D. A. Galbraith, X. Yang, E. L. Niño, S. Yi, and C. Grozinger. Parallel Epigenomic and Transcriptomic Responses to Viral Infection in Honey Bees (Apis mellifera). PLoS Pathogens, 11(3):1–24, 2015. ISSN 15537374. doi: 10.1371/journal.ppat.1004713.

L. Y. Geer, A. Marchler-Bauer, R. C. Geer, L. Han, J. He, S. He, C. Liu, W. Shi, and S. H. Bryant. The NCBI BioSystems database. Nucleic Acids Research, 38(SUPPL.1):492–496, 2009. ISSN 03051048. doi: 10.1093/nar/gkp858.

G. Harwood, G. Amdam, and D. Freitak. The role of vitellogenin in the transfer of immune elicitors from gut to hypopharyngeal glands in honey bees (apis mellifera). Journal of Insect Physiology, 112:90–100, 2019. ISSN 0022-1910. doi: https://doi.org/10.1016/j.jinsphys.2018.12.006. URL https://www.sciencedirect.com/science/article/pii/S0022191018302300.

H. Havukainen, D. Münch, A. Baumann, S. Zhong, Ø. Halskau, M. Krogsgaard, and G. V. Amdam. Vitellogenin recognizes cell damage through membrane binding and shields living cells from reactive oxygen species. Journal of Biological Chemistry, 288(39):28369–28381, 2013.

A. Hayward, T. Takahashi, W. G. Bendena, S. S. Tobe, and J. H. Hui. Comparative genomic and phylogenetic analysis of vitellogenin and other large lipid transfer proteins in metazoans. FEBS letters, 584(6):1273–1278, 2010.

W. Hennig. Insect Phylogeny. Wiley-Blackwell, 1981.

J. F. Hillyer. Insect immunology and hematopoiesis. Developmental and Comparative Immunology, 58:102–118, 2016. ISSN 18790089. doi: 10.1016/j.dci.2015.12.006.

S. Holm. A simple sequentially rejective multiple test procedure. Scandinavian journal of statistics, pages 65–70, 1979.

A. C. A. Hope. A simplified monte carlo significance test procedure. Journal of the Royal Statistical Society. Series B (Methodological), 30(3):582–598, 1968. ISSN 00359246. URL http://www.jstor.org/stable/2984263.

J. Huerta-Cepas, D. Szklarczyk, K. Forslund, H. Cook, D. Heller, M. C. Walter, T. Rattei, D. R. Mende, S. Sunagawa, M. Kuhn, L. J. Jensen, C. Von Mering, and P. Bork. EGGNOG 4.5: A hierarchical orthology framework with improved functional annotations for eukaryotic, prokaryotic and viral sequences. Nucleic Acids Research, 44(D1):D286–D293, 2016. ISSN 13624962. doi: 10.1093/nar/gkv1248.

J. Huerta-Cepas, K. Forslund, L. P. Coelho, D. Szklarczyk, L. J. Jensen, C. Von Mering, and P. Bork. Fast genome-wide functional annotation through orthology assignment by eggNOG-mapper. Molecular Biology and Evolution, 34(8):2115–2122, 2017. ISSN 15371719. doi: 10.1093/molbev/msx148.

K. J. Ishii, C. Coban, H. Kato, K. Takahashi, Y. Torii, F. Takeshita, H. Ludwig, G. Sutter, K. Suzuki, H. Hemmi, S. Sato, M. Yamamoto, S. Uematsu, T. Kawai, O. Takeuchi, and S. Akira. A toll-like receptor– independent antiviral response induced by double-stranded b-form dna. Nature Immunology, 7(1):40–48, 2006. doi: 10.1038/ni1282. URL https://doi.org/10.1038/ni1282.

K. M. Kapheim, H. Pan, C. Li, S. L. Salzberg, D. Puiu, T. Magoc, H. M. Robertson, M. E. Hudson, and A. Venkat. Genomic signatures of evolutionary transitions from solitary to group living. Science, 348(6239):1139–1143, 2015. ISSN 0036-8075. doi: 10.1126/science.aaa4788.

K. Katoh, J. Rozewicki, and K. D. Yamada. MAFFT online service: multiple sequence alignment, interactive sequence choice and visualization. Briefings in Bioinformatics, (July): 1–7, 2017. ISSN 1467-5463. doi: 10.1093/bib/bbx108. URL http://academic.oup.com/bib/article/doi/10.1093/bib/bbx108/4106928/MAFFT-online-service-multiple-sequence-alignment.

S. D. Kocher, C. Li, W. Yang, H. Tan, S. V. Yi, X. Yang, H. E. Hoekstra, G. Zhang, N. E. Pierce, and D. W. Yu. The draft genome of a socially polymorphic halictid. Genome Biology, 14:R142, 2013. ISSN 1474760X. doi: 10.1186/s13059-014-0574-0.

C.-J. Kuo, M. Hansen, and E. Troemel. Autophagy and innate immunity: Insights from invertebrate model organisms. Autophagy, 14(2):233–242, 2018. doi: 10.1080/15548627.2017.1389824. URL https://doi.org/10.1080/15548627.2017.1389824. PMID: 29130360.

G. Liu, Z. Li, M. Yang, L. Lin, J. Liu, and M. Chen. Functional characterization of a putative lipopolysaccharide-induced tnf-alpha factor (litaf) from blood clam tegillarca granosa in innate immunity. Fish & Shellfish Immunology, 97:390–402, 2020. ISSN 1050-4648. doi: https://doi.org/10.1016/j.fsi.2019.12.051. URL https://www.sciencedirect.com/science/article/pii/S1050464819311805.

A. P. Lourenço, J. R. Martins, M. M. Bitondi, and Z. L. Simões. Trade-off between immune stimulation and expression of storage protein genes. Archives of Insect Biochemistry and Physiology: Published in Collaboration with the Entomological Society of America, 71(2):70–87, 2009.

A. P. Lourenço, M. M. Florecki, Z. L. P. Simões, and J. D. Evans. Silencing of apis mellifera dorsal genes reveals their role in expression of the antimicrobial peptide defensin-1. Insect molecular biology, 27(5):577–589, 2018.

F. Manfredini, D. Shoemaker, and C. M. Grozinger. Dynamic changes in host– virus interactions associated with colony founding and social environment in fire ant queens (solenopsis invicta). Ecology and evolution, 6(1):233–244, 2016.

A. McMenamin, K. Daughenbaugh, F. Parekh, M. Pizzorno, and M. Flenniken. Honey Bee and Bumble Bee Antiviral Defense. Viruses, 10(8):395, 2018. ISSN 1999-4915. doi: 10.3390/v10080395. URL http://www.mdpi.com/1999-4915/10/8/395.

F. K. Mendes, D. Vanderpool, B. Fulton, and M. W. Hahn. CAFE 5 models variation in evolutionary rates among gene families. Bioinformatics, Dec. 2020.

C. Mendoza, P. Olguın, G. Lafferte, U. Thomas, S. Ebitsch, E. D. Gundelfinger, M. Kukuljan, and J. Sierralta. Novel isoforms of dlg are fundamental for neuronal development indrosophila. Journal of Neuroscience, 23(6):2093– 2101, 2003.

B. Milutinović, R. Peuß, K. Ferro, and J. Kurtz. Immune priming in arthropods: an update focusing on the red flour beetle. Zoology, 119(4):254–261, 2016.

M. Navajas, A. Migeon, C. Alaux, M. Martin-Magniette, G. Robinson, J. Evans, S. Cros-Arteil, D. Crauser, and Y. Le Conte. Differential gene expression of the honey bee Apis mellifera associated with Varroa destructor infection. BMC Genomics, 9(1):301, 2008. ISSN 1471-2164. doi: 10.1186/1471-2164-9-301. URL http://bmcgenomics.biomedcentral.com/articles/10.1186/1471-2164-9-301.

NCBI Resource Coordinators. Database resources of the national center for biotechnology information. Nucleic acids research, 46(D1):D8–D13, 01 2018. doi: 10.1093/nar/gkx1095. URL https://pubmed.ncbi.nlm.nih.gov/29140470.

D. J. Obbard, F. M. Jiggins, D. L. Halligan, and T. J. Little. Natural selection drives extremely rapid evolution in antiviral RNAi genes. Current Biology, 16(6):580–585, 2006. ISSN 09609822. doi: 10.1016/j.cub.2006.01.065.

D. J. Obbard, K. H. J. Gordon, A. H. Buck, and F. M. Jiggins. The evolution of rnai as a defence against viruses and transposable elements. Philos Trans R Soc Lond B Biol Sci, 364(1513):99–115, Jan 2009. ISSN 1471-2970 (Electronic); 0962-8436 (Print); 0962-8436 (Linking). doi: 10.1098/rstb.2008.0168.

S. Otani, N. Bos, and S. H. Yek. Transitional Complexity of Social Insect Immunity. Frontiers in Ecology and Evolution, 4(69):doi: 10.3389/fevo.2016.00069, 2016. ISSN 2296-701X. doi: 10.3389/fevo.2016.00069. URL http://journal.frontiersin.org/Article/10.3389/fevo.2016.00069/abstract.

P. R. Oxley, M. Spivak, and B. P. Oldroyd. Six quantitative trait loci influence task thresholds for hygienic behaviour in honeybees (Apis mellifera). Molecular Ecology, 19(7):1452–1461, 2010. ISSN 09621083. doi: 10.1111/j.1365-294X.2010.04569.x.

E.-M. Park, Y.-O. Kim, B.-H. Nam, H. J. Kong, W.-J. Kim, S.-J. Lee, I.-S. Kong, and T.-J. Choi. Cloning, characterization and expression analysis of the gene for a putative lipopolysaccharide-induced tnf-*α* factor of the pacific oyster, crassostrea gigas. Fish & Shellfish Immunology, 24 (1):11–17, 2008. ISSN 1050-4648. doi: https://doi.org/10.1016/j.fsi.2007.07.003. URL https://www.sciencedirect.com/science/article/pii/S1050464807001234.

H. G. Park, K. S. Lee, B. Y. Kim, H. J. Yoon, Y. S. Choi, K. Y. Lee, H. Wan, J. Li, and B. R. Jin. Honeybee (apis cerana) vitellogenin acts as an antimicrobial and antioxidant agent in the body and venom. Developmental & Comparative Immunology, 85:51–60, 2018. ISSN 0145-305X. doi: https://doi.org/10.1016/j.dci.2018.04.001. URL https://www.sciencedirect.com/science/article/pii/S0145305X18300958.

R Core Team. *R: A Language and Environment for Statistical Computing*. R Foundation for Statistical Computing, Vienna, Austria, 2020. URL https://www.R-project.org/.

S. M. Rehan and A. L. Toth. Climbing the social ladder: The molecular evolution of sociality. Trends in Ecology and Evolution, 30(7):426–433, 2015. ISSN 01695347. doi: 10.1016/j.tree.2015.05.004. URL http://dx.doi.org/10.1016/j.tree.2015.05.004.

S. M. Rehan, K. M. Glastad, S. P. Lawson, and B. G. Hunt. The Genome and Methylome of a Subsocial Small Carpenter Bee, Ceratina calcarata. Genome biology and evolution, 8(5):1401–1410, 2016. ISSN 17596653. doi: 10.1093/gbe/evw079.

F.-J. Richard, H. L. Holt, and C. M. Grozinger. Effects of immunostimulation on social behavior, chemical communication and genome-wide gene expression in honey bee workers (Apis mellifera). BMC Genomics, 13(1):558, 2012. ISSN 1471-2164. doi: 10.1186/1471-2164-13-558. URL http://bmcgenomics.biomedcentral.com/articles/10.1186/1471-2164-13-558.

B. E. Rubin, B. M. Jones, B. Hunt, and S. D. Kocher. Rate variation in the evolution of non-coding DNA associated with social evolution in bees. Phil. Trans. R. Soc. B, 374:20180247, 2019. doi: 10.1101/461079. URL https://www.biorxiv.org/content/10.1101/461079v2.

T. B. Sackton. Comparative genomics and transcriptomics of host–pathogen interactions in insects: evolutionary insights and future directions. Current Opinion in Insect Science, 31:106–113, 2019. ISSN 22145753. doi: 10.1016/j.cois.2018.12.007. URL https://doi.org/10.1016/j.cois.2018.12.007.

T. B. Sackton. Studying Natural Selection in the Era of Ubiquitous Genomes. Trends in Genetics, pages 1–12, 2020. ISSN 13624555. doi: 10.1016/j.tig.2020.07.008. URL https://doi.org/10.1016/j.tig.2020.07.008.

T. B. Sackton, B. P. Lazzaro, T. A. Schlenke, J. D. Evans, D. Hultmark, and A. G. Clark. Dynamic evolution of the innate immune system in drosophila. Nature Genetics, 39(12):1461–1468, 2007. doi: 10.1038/ng.2007.60. URL https://doi.org/10.1038/ng.2007.60.

T. B. Sackton, J. H. Werren, and A. G. Clark. Characterizing the infection-induced transcriptome of Nasonia vitripennis reveals a preponderance of taxonomically-restricted immune genes. PLoS ONE, 8(12):e83984, 2013. ISSN 19326203. doi: 10.1371/journal.pone.0083984.

B. M. Sadd, Y. Kleinlogel, R. Schmid-Hempel, and P. Schmid-Hempel. Trans-generational immune priming in a social insect. Biology Letters, 1(4):386– 388, 2005. ISSN 1744-9561. doi: 10.1098/rsbl.2005.0369. URL http://rsbl.royalsocietypublishing.org/cgi/doi/10.1098/rsbl.2005.0369.

B. M. Sadd, S. M. Barribeau, G. Bloch, D. C. de Graaf, P. Dearden, C. G. Elsik, J. Gadau, C. J. Grimmelikhuijzen, M. Hasselmann, J. D. Lozier, H. M. Robertson, G. Smagghe, E. Stolle, M. Van Vaerenbergh, R. M. Waterhouse, E. Bornberg-Bauer, S. Klasberg, A. K. Bennett, F. Câmara, R. Guigó, K. Hoff, M. Mariotti, M. Munoz-Torres, T. Murphy, D. Santesmasses, G. V. Amdam, M. Beckers, M. Beye, M. Biewer, M. M. Bitondi, M. L. Blaxter, A. F. Bourke, M. J. Brown, S. D. Buechel, R. Cameron, K. Cappelle, J. C. Carolan, O. Christiaens, K. L. Ciborowski, D. F. Clarke, T. J. Colgan, D. H. Collins, A. G. Cridge, T. Dalmay, S. Dreier, L. du Plessis, E. Duncan, Erler, J. Evans, T. Falcon, K. Flores, F. C. Freitas, T. Fuchikawa, S. Gempe, K. Hartfelder, F. Hauser, S. Helbing, F. C. Humann, F. Irvine, L. S. Jermiin, C. E. Johnson, R. M. Johnson, A. K. Jones, T. Kadowaki, J. H. Kidner, V. Koch, A. Köhler, F. B. Kraus, H. M. G. Lattorff, M. Leask, G. A. Lockett, E. B. Mallon, D. S. M. Antonio, M. Marxer, I. Meeus, R. F. Moritz, A. Nair, K. Näpflin, I. Nissen, J. Niu, F. M. Nunes, J. G. Oakeshott, A. Osborne, M. Otte, D. G. Pinheiro, N. Rossié, O. Rueppell, C. G. Santos, R. Schmid-Hempel, B. D. Schmitt, C. Schulte, Z. L. Simões, M. P. Soares, L. Swevers, E. C. Winnebeck, F. Wolschin, N. Yu, E. M. Zdobnov, P. K. Aqrawi, K. P. Blankenburg, M. Coyle, L. Francisco, A. G. Hernandez, M. Holder, M. E. Hudson, L. Jackson, J. Jayaseelan, V. Joshi, C. Kovar, S. L. Lee, R. Mata, T. Mathew, I. F. Newsham, R. Ngo, G. Okwuonu, C. Pham, L.-L. Pu, N. Saada, J. Santibanez, D. Simmons, R. Thornton, A. Venkat, K. K. Walden, Y.-Q. Wu, G. Debyser, B. Devreese, C. Asher, J. Blommaert, A. D. Chipman, L. Chittka, B. Fouks, J. Liu, M. P. O’Neill, S. Sumner, D. Puiu, J. Qu, S. L. Salzberg, S. E. Scherer, D. M. Muzny, S. Richards, G. E. Robinson, R. A. Gibbs, P. Schmid-Hempel, and K. C. Worley. The genomes of two key bumblebee species with primitive eusocial organization. Genome Biology, 16(1):76, 2015. ISSN 1465-6906. doi: 10.1186/s13059-015-0623-3. URL http://genomebiology.com/2015/16/1/76.

H. Salmela and L. Sundström. Vitellogenin in inflammation and immunity in social insects. Inflammation and Cell Signaling, 5, 2018.

H. Salmela, G. V. Amdam, and D. Freitak. Transfer of immunity from mother to offspring is mediated via egg-yolk protein vitellogenin. PLOS Pathogens, 11(7):1–12, 07 2015. doi: 10.1371/journal.ppat.1005015. URL https://doi.org/10.1371/journal.ppat.1005015.

T. A. Schlenke, J. Morales, S. Govind, and A. G. Clark. Contrasting infection strategies in generalist and specialist wasp parasitoids of drosophila melanogaster. PLoS Pathog, 3(10):e158, 2007.

S.-C. Seehuus, K. Norberg, U. Gimsa, T. Krekling, and G. V. Amdam. Reproductive protein protects functionally sterile honey bee workers from oxidative stress. Proceedings of the National Academy of Sciences, 103(4): 962–967, 2006.

G. Sheehan, A. Garvey, M. Croke, and K. Kavanagh. Innate humoral immune defences in mammals and insects: The same, with differences? Virulence, 9(1):1625–1639, 2018.

W. A. Shell, M. A. Steffen, H. K. Pare, A. S. Seetharam, A. J. Severin, A. L. Toth, and S. M. Rehan. Sociality sculpts similar patterns of molecular evolution in two independently evolved lineages of eusocial bees. Commun Biol, 4(1):253, Feb. 2021.

N. Silverman and T. Maniatis. Nf-*κ*b signaling pathways in mammalian and insect innate immunity. Genes & development, 15(18):2321–2342, 2001.

R. C. Smith, A. G. Eappen, A. J. Radtke, and M. Jacobs-Lorena. Regulation of anti-plasmodium immunity by a litaf-like transcription factor in the malaria vector anopheles gambiae. PLoS pathogens, 8(10):e1002965, 2012. ISSN 1553-7366. doi: 10.1371/journal.ppat.1002965. URL https://europepmc.org/articles/PMC3475675.

F. Supek, M. Bǒsnjak, N. Škunca, and T. Šmuc. Revigo summarizes and visualizes long lists of gene ontology terms. PLoS ONE, 6(7), 2011. ISSN 19326203. doi: 10.1371/journal.pone.0021800.

M. Suyama, D. Torrents, P. Bork, and M. Delbru. PAL2NAL : robust conversion of protein sequence alignments into the corresponding codon alignments. Nucleic Acids Research, 34:609–612, 2006. doi: 10.1093/nar/gkl315.

X. Tang, D. L. Marciano, S. E. Leeman, and S. Amar. Lps induces the interaction of a transcription factor, lps-induced tnf-*α* factor, and stat6(b) with effects on multiple cytokines. Proceedings of the National Academy of Sciences, 102(14):5132–5137, 2005. ISSN 0027-8424. doi: 10.1073/pnas.0501159102. URL https://www.pnas.org/content/102/14/5132.

J. Trapp, A. McAfee, and L. J. Foster. Genomics, transcriptomics and proteomics: enabling insights into social evolution and disease challenges for managed and wild bees. Molecular Ecology, 26(3):718–739, 2017. ISSN 1365294X. doi: 10.1111/mec.13986.

K. Troha, J. H. Im, J. Revah, B. P. Lazzaro, and N. Buchon. Comparative transcriptomics reveals CrebA as a novel regulator of infection tolerance in D. melanogaster, volume 14. 2018. ISBN 1111111111. doi: 10.1371/journal.ppat.1006847.

A. Wallberg, I. Bunikis, O. V. Pettersson, M. B. Mosbech, A. K. Childers, J. D. Evans, A. S. Mikheyev, H. M. Robertson, G. E. Robinson, and M. T. Webster. A hybrid de novo genome assembly of the honeybee, Apis mellifera, with chromosome-length scaffolds. BMC Genomics, 20(1):1–19, 2019. ISSN 14712164. doi: 10.1186/s12864-019-5642-0.

P.-H. Wang, D.-H. Wan, L.-R. Pang, Z.-H. Gu, W. Qiu, S.-P. Weng, X.-Q. Yu, and J.-G. He. Molecular cloning, characterization and expression analysis of the tumor necrosis factor (tnf) superfamily gene, tnf receptor superfamily gene and lipopolysaccharide-induced tnf-*α* factor (litaf) gene from litopenaeus vannamei. Developmental & Comparative Immunology, 36 (1):39–50, 2012. ISSN 0145-305X. doi: https://doi.org/10.1016/j.dci.2011.06.002. URL https://www.sciencedirect.com/science/article/pii/S0145305X11001686.

Q. Wang, M. Ren, X. Liu, H. Xia, and K. Chen. Peptidoglycan recognition proteins in insect immunity. Molecular Immunology, 106:69–76, 2019. ISSN 0161-5890. doi: https://doi.org/10.1016/j.molimm.2018.12.021. URL https://www.sciencedirect.com/science/article/pii/S0161589018309015.

E. O. Wilson and B. Hölldobler. Eusociality: origin and consequences. Proceedings of the National Academy of Sciences, 102(38):13367–13371, 2005.

N. Wilson-Rich, M. Spivak, N. H. Fefferman, and P. T. Starks. Genetic, Individual, and Group Facilitation of Disease Resistance in Insect Societies. Annual Review of Entomology, 54(1):405–423, 2009. ISSN 0066-4170. doi: 10.1146/annurev.ento.53.103106.093301. URL http://www.annualreviews.org/doi/10.1146/annurev.ento.53.103106.093301.

G. S. Withers, N. F. Day, E. F. Talbot, H. E. Dobson, and C. S. Wallace. Experience-dependent plasticity in the mushroom bodies of the solitary bee osmia lignaria (megachilidae). Developmental neurobiology, 68(1):73–82, 2008.

D. F. Woods and P. J. Bryant. The discs-large tumor suppressor gene of drosophila encodes a guanylate kinase homolog localized at septate junctions. Cell, 66(3):451–464, 1991.

X.-P. Xiong, K. Kurthkoti, K.-Y. Chang, J.-L. Li, X. Ren, J.-Q. Ni, T. M. Rana, and R. Zhou. mir-34 modulates innate immunity and ecdysone signaling in drosophila. PLoS pathogens, 12(11):e1006034, 2016.

Y. Xu. Dna damage: a trigger of innate immunity but a requirement for adaptive immune homeostasis. Nature Reviews Immunology, 6(4):261–270, 2006. doi: 10.1038/nri1804. URL https://doi.org/10.1038/nri1804.

Z. Yang. PAML: a program package for phylogenetic analysis by maximum likelihood. Computer applications in the biosciences : CABIOS, 13(5):555– 6, 1997. ISSN 0266-7061. URL http://www.ncbi.nlm.nih.gov/pubmed/9367129.

Z. Yang. PAML 4: Phylogenetic analysis by maximum likelihood. Molecular Biology and Evolution, 24(8):1586–1591, 2007. ISSN 07374038. doi: 10.1093/molbev/msm088.

Z. Yang and R. Nielsen. Codon-Substitution Models for Detecting Molecular Adaptation at Individual Sites Along Specific Lineages. Mol. Biol. Evol., 19(6):908–917, 2002.

E. M. Zdobnov, F. Tegenfeldt, D. Kuznetsov, R. M. Waterhouse, F. A. Simao, P. Ioannidis, M. Seppey, A. Loetscher, and E. V. Kriventseva. OrthoDB v9.1: Cataloging evolutionary and functional annotations for animal, fungal, plant, archaeal, bacterial and viral orthologs. Nucleic Acids Research, 45 (D1):D744–D749, 2017. ISSN 13624962. doi: 10.1093/nar/gkw1119.

D. Žgur-Bertok. Dna damage repair and bacterial pathogens. PLoS pathogens, 9(11):e1003711–e1003711, 2013. doi: 10.1371/journal.ppat.1003711. URL https://pubmed.ncbi.nlm.nih.gov/24244154.

J. Zhang, R. Nielsen, and Z. Yang. Evaluation of an Improved Branch-Site Likelihood Method for Detecting Positive Selection at the Molecular Level. Mol. Biol. Evol., 22(12):2472–2479, 2005. doi: 10.1093/molbev/msi237.

S. Zhang, S. Wang, H. Li, and L. Li. Vitellogenin, a multivalent sensor and an antimicrobial effector. The international journal of biochemistry & cell biology, 43(3):303–305, 2011.

J. Zou, P. Guo, N. Lv, and D. Huang. Lipopolysaccharide-induced tumor necrosis factor-*α*factor enhances inflammation and is associated with cancer (review). Mol Med Rep, 12(5):6399–6404, 2015. doi: 10.3892/mmr.2015.4243. URL https://doi.org/10.3892/mmr.2015.4243.

